# Iridescent structural coloration in a crested Cretaceous enantiornithine bird from Jehol Biota

**DOI:** 10.1101/2024.10.21.619362

**Authors:** Zhiheng Li, Jinsheng Hu, Thomas A. Stidham, Mao Ye, Min Wang, Yanhong Pan, Tao Zhao, Jingshu Li, Zhonghe Zhou, Julia A. Clarke

## Abstract

A combination of sectioning and microscopy techniques, along with the application of finite-difference-time-domain modeling on a fossil feather, results in the novel estimation of the range of iridescent colors from the fossilized melanosome type and organization preserved in the elongate head crest feathers of a new Cretaceous enantiornithine bird. The densely packed rod-like melanosomes are estimated to have yielded from red to deep blue iridescent coloration of the head feathers. The shape and density of these melanosomes also may have further increased the feather’s structural strength. This occurrence on a likely male individual is highly suggestive of both a signaling function of the iridescent crest, and a potential behavioral role in adjusting the angle of light incidence to control the display of this iridescent structural coloration.

## Introduction

The recognition and study of melanosomes in fossil feathers of non-avialan dinosaurs and birds has added greater complexity to the pattern of their diversity and macroevolution, demonstrating the intertwined evolution of coloration with feather functions across display, thermoregulation, and other uses (Li, Gao et al. 2012, Li, Clarke et al. 2014, Terrill and Shultz 2023). This linkage among colors, structures, and functions early in feather evolution extends even to convergence in iridescent feathers among non-avialan theropods, and it is highly suggestive of commonalities shared among stem and crown birds in terms of their use of colored pigments for camouflage, structural strength, and display (Hu, Clarke et al. 2018, Foth and Rauhut 2020). In a comparative setting with the melanosome feather coloration of living birds, workers have endeavored to reconstruct the colors of fossil feathers using quantitative means focused predominantly on melanosome geometry and density, and the results point to the fossilized melanosomes having produced patterns of black, brown, rufous, grey, and other colors (Vinther, Briggs et al. 2008, Li, Gao et al. 2010, Zhang, Kearns et al. 2010, Li, Gao et al. 2012, Peteya, Clarke et al. 2017, Benton, Dhouailly et al. 2019). Furthermore, the specific shape and arrangement of some fossilized feather melanosomes derived from both non-avialan theropod dinosaurs and birds are similar to those required for producing structurally-based iridescent feather coloration in living birds, a case of independent evolution (Stoddard and Prum 2011, Hu, Clarke et al. 2018, Pan, Li et al. 2022).

Feathers serve multiple functions in birds from comprising an insulatory epidermal layer to forming aerodynamic flight surfaces, and as structures for the containment and display of various pigments. Living birds exhibit diverse feather morphologies and colors, and the significant diversity of known fossil feathers, along with their preserved pigments, points to their past use for camouflage, as well as display and signaling (Prum 2006, Foth and Rauhut 2020, Terrill and Shultz 2023). While the display of most colored feathers are passive and available to any viewer, birds can behaviorally control their signaling through: the erection of feathered ornaments such as the head crest in hoopoes (*Upupa*) or Peacocks (*Pavo*) tails; the exposure of normally hidden brightly colored crown feathers as in Kinglets (*Regulus*); and even the refined control of the black and iridescent coloration in birds such as hummingbirds (Trochilidae) and birds of paradise (Paradisaeidae) (Zi, Yu et al. 2003, Wilts, Michielsen et al. 2014, Stoddard, Eyster et al. 2020). This great variation in structural or iridescent colors results from differences in pigments, their organization, and other structural modifications. For example, the variety of mixed structural colors present in a peacock feather are achieved through changes in the spacing in the keratinous matrix and melanosome organization (Zi, Yu et al. 2003). In addition to the normal range of spectral coloration, hummingbirds can see non-spectral colors as well, and their utility is supported by vision perception experiments (Stoddard, Eyster et al. 2020). The interplay of bird of paradise feather coloration and their vision similarly indicates coordination of their mating displays with spectral properties through the avian visual system (Wilts, Michielsen et al. 2014). Individual birds actively control the signaling functions of their feathers with feathers or ornaments hidden/displayed or even the color presented (as dark/black vs. vibrant/iridescent) being consciously restricted or directed to particular viewers such as potential mates or rivals (Wilts, Michielsen et al. 2014).

Fossilized feathers typically have been evaluated within two-dimensional surface views, limiting the understanding of their overall ultrastructure and perhaps their true coloration. As a step forward in our reconstruction of the structure and function (color display) of early bird feathers, we examined the unique head-crest feathers of a well-preserved specimen (Institute of Vertebrate Paleontology and Paleoanthropology, IVPP V26899) of the Early Cretaceous (∼120 Ma) enantiornithine *Shangyang* sp. from the Jiufotang Formation of northeastern China. This fossil preserves a diverse feather assemblage across the body with dense contour feathers, flight remiges, and two rachis-dominated tail feathers (Figure 1 and figure S1). Among its diverse feather morphologies, this specimen also displays a strikingly enlarged feathered crest on the head that uniquely extends from the caudal end of the skull to a position rostral to the nasal bone (Figure 1). For our study, we sampled a long isolated mature feather that is displaced from the caudal part of the crest, as well as three feather fragments extracted from the *in-situ* position of the head crest.

**Figure 1.**
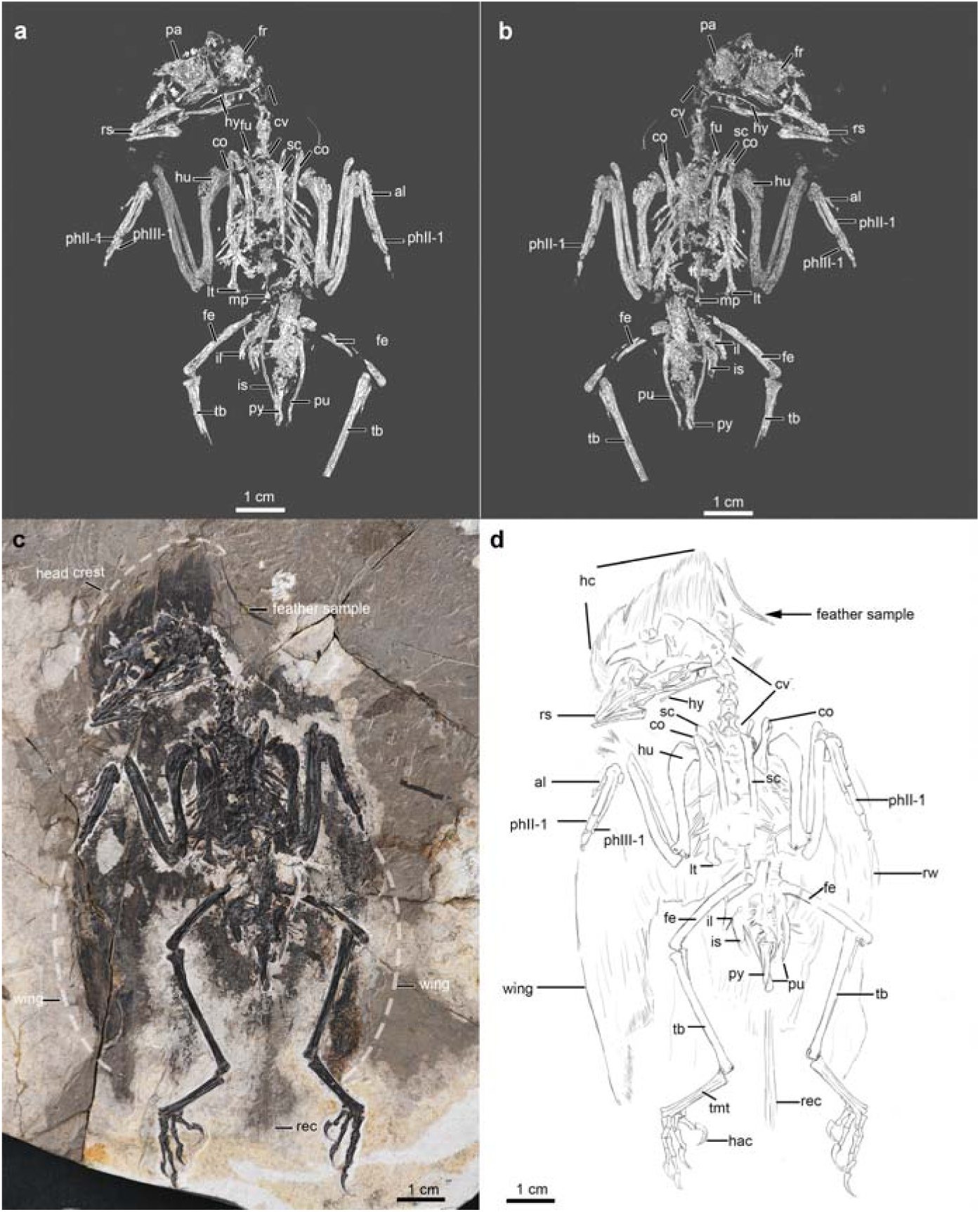
Digital rendering, photography, and line-drawing of the enantiornithine fossil specimen (*Shanyang* sp., IVPP V26899) (a), (c), (d) - dorsal; (b)- ventral view showing the major skeletal elements. The feather extracted from the crown are labeled as feather sample (**c**). Abbreviations: al, alular metacarpal; co, coracoid; cv, cervical vertebra; fe, femur; fi, fibula; fr frontal; fu, furcula; hac, hallucal ungual; hc, head crest; hu, humerus; hy, hyoid; il, ilium; is, ischium; lt, lateral trabecula; mcf, molted crest feather; mc II, major metacarpal; mp, midline process; ph II-1, first digit of phalanx II; phIII-1, first digit of phalanx III; pa, parietal; pu, pubis; py, pygostyle; r, radius; rec, rectrices; rem, remiges; rs, rostrum; rw, right wing; sc, scapula; sk, skull; st, sternum; tb, tibiotarsus; and tmt, tarsometatarsus.

In this study, we utilized serial ultrathin histological sectioning, scanning/transmission electron microscopy (S/TEM) imaging, and finite-difference time-domain (FDTD) modeling to investigate the potential iridescent coloration mechanisms in the elongated crest feathers from this newly discovered enantiornithine specimen of the Jehol Biota. Differing from previous statistical reconstructions of fossil color that measured melanosome aspect ratios, our new approach takes the three-dimensional shape and packing patterns of melanosomes, as well as the keratinous matrix, into consideration. With these novel modeling approaches, a more accurate coloration estimation was produced for its crest feather.

## Results

In the new fossil, head crest feathers were selected from the caudal side of the head and sectioned using ultrathin serial sectioning. The overall shape of the sampled crest feather is well-preserved and comprised of elongate and relatively wide barbs. The dominant barb length is proportionally longer than the feather barb examined from the neck and head region of *Confuciusornis* (Foth 2012). The feather is different from the bilaterally symmetric plumulaceous feathers of living birds, and we define it as a barb-dominant feather, since most of the exposed feather tissue is composed of long barbs with a very short and thick calamus. It is estimated to correlate with evolutionary stage II and IIIa of (Prum and Brush 2002), similar to the intermediate feather type proposed as found in an amber fossil (Perrichot, Marion et al. 2008). The new feather described here is distinct from the amber feather in showing an asymmetric shape, radically diverging toward one side (caudal side), and is much larger in size (Perrichot, Marion et al. 2008). There are dense clusters inside the barbs as seen from images of both SEM and TEM.

Ultrathin histological sectioning and application of SEM and S/TEM imaging of the fossil feather samples (see Supplementary materials) resulted in the identification of partially interlinked melanosomes with a derived asymmetric packing pattern (Figures 2 and 3). The long axial crack in Figure 2 may represent the surface of the barbs, resembling a crack formed in maturation experiments (Zhao, Hu et al. 2020). Those melanosome clusters are present and concentrated mostly within the barbs rather than in the barbules of extant bird feathers based on similar size (Figure 2). The long axis of the densely packed melanosomes is aligned overall parallel with the barb’s long axis (Figure 2), which has been confirmed by detailed histological data as well. The melanosomes are arranged in an asymmetric hexagonal conformation, clearly visible in the different sections (see Figure 3).

**Figure 2.**
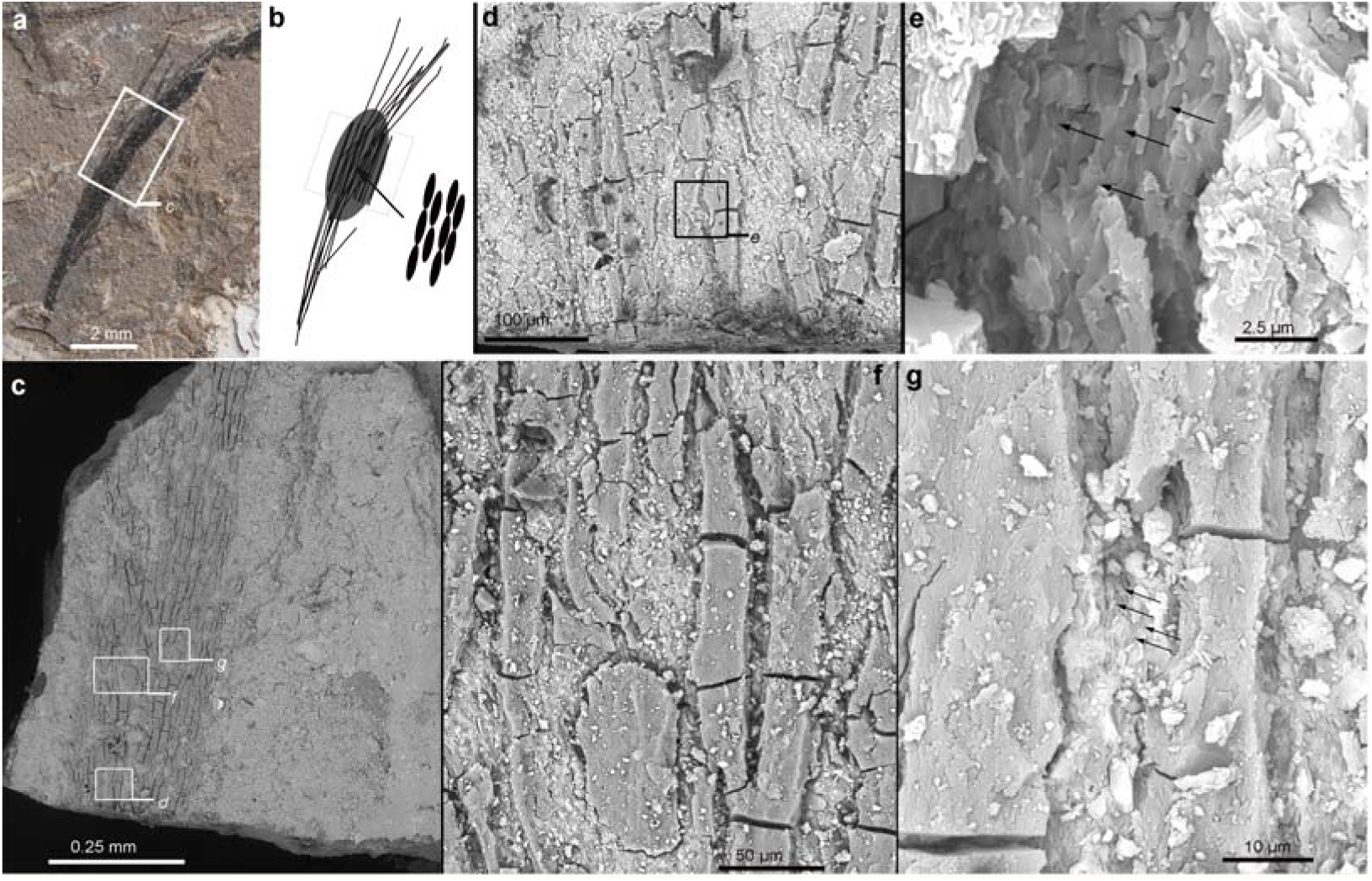
Photograph (a) and simplified line-diagram (b) of the sampled isolated long feather with a focused area from SEM imaging (c-g). As shown in the SEM images, the long elliptical or oval-shaped melanosomes are tightly aligned with their long axis parallel to the elongated barbs **(e, g**). The position of enlarged images (**d, e, f, and g**) is indicated as square in (**c, d**); black arrows in (**e**) and (**g**) indicate the melanosome clusters.

**Figure 3.**
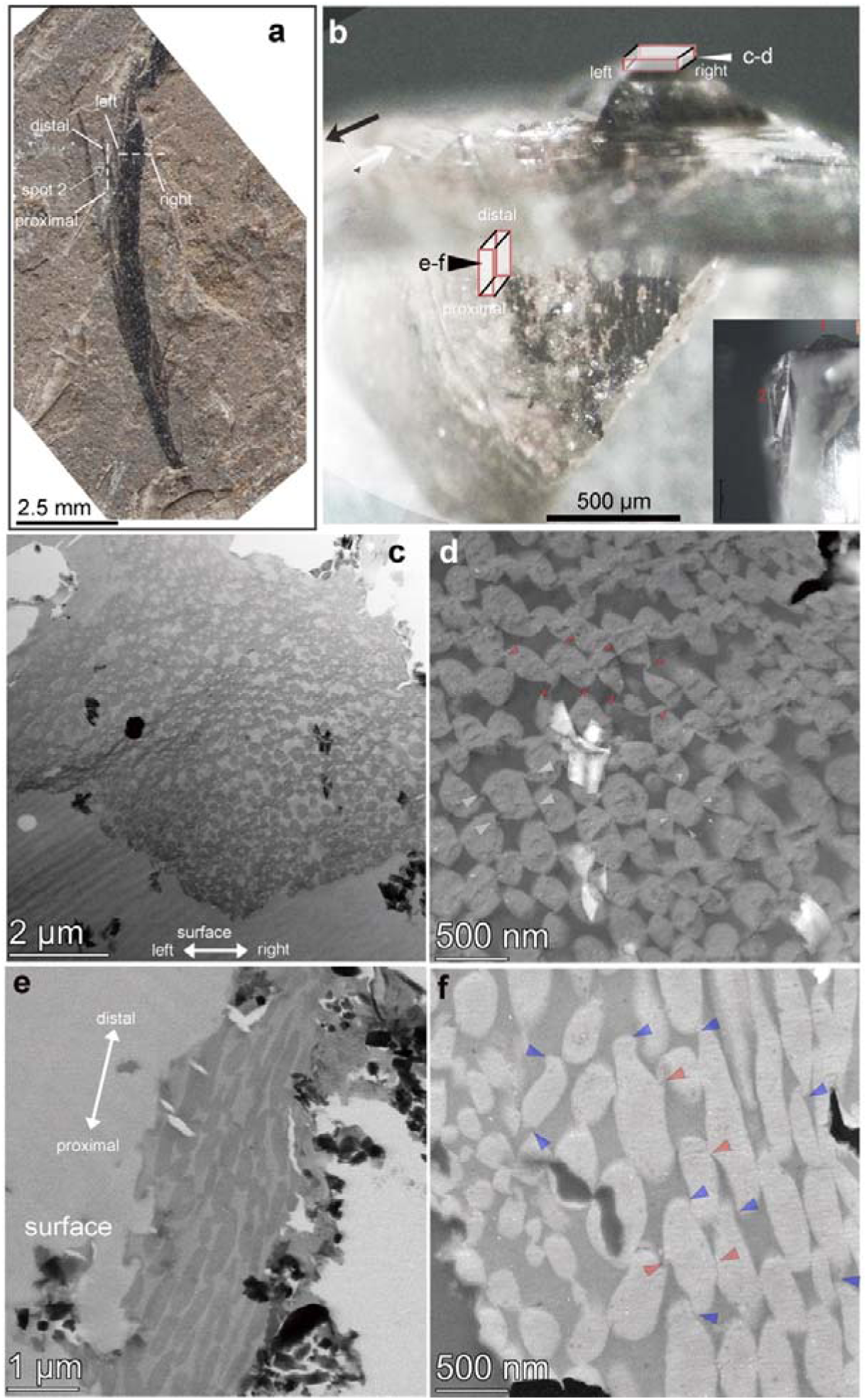
Photograph of the crest feather (a) and prepared blocks of the feather barb (b) for thin sectioning. The sampled spots for the sliced feather are indicated as long red-colored cubes with their respective positions in (**b**). (**c**) to (**f**) are (STEM) images obtained from individual slices as indicated in (**b**). The shape and arrangement of the melanosome packing (ACHP, asymmetric compact hexagonal packing) is clearly revealed in (**d**); (**c**) and (**d**) are derived from the cross-section; (**e**) and (**f**) are derived from the nearly longitudinal section. As indicated by the red arrows in (d) and (f), the dorsal and ventral hooks are dominant in the cross-sectional images. On the other hand, hooks connecting the proximal and distal arrays of melanosomes are better visualized in the longitudinal section, as labeled with blue arrows in (f). Orientation of the slices were labeled (left, right, proximal, distal) with their respective petitioning in the fossil before and during sectioning.

SEM imaging of the cross section and surface view of the sampled feather indicate the presence of a few major barbs without barbules (figures S1 and S3-4). The thickness of the complete feather barb with filled melanosomes (preserved) is around 10 µm, which is much greater than the typical diameter of known barbules, but similar to barbs in extant birds (Freyer, Wilts et al. 2021). The three-dimensionally packed melanosomes were studied in both cross and nearly longitudinal sections within the dark colored barbs (one in the middle of the barbs and another in the lateral side of the feather block (Figure 3a, b).

Hooks are commonly observed on the oval-shaped melanosomes in cross-sectional views, with two dominant types identified on the dorsal and ventral sides (Figure 3c-d, red arrows). These hooks are deflected in opposing directions, linking melanosomes from different arrays (dorsal-ventral) together. The major axis(y) of the oval-shaped melanosomes (mean = 283 nm) is oriented toward the left side in cross-section, while the shorter axis(x) measures approximately 186 nm (Table S2). In oblique or near-longitudinal sections (Figure 3e-f), the hooked structures’ connections to the distal and proximal sides of neighboring melanosomes are clearly visible (blue arrows, Figure 3f). The mean long axis (z) of the melanosomes is approximately 1774 nm (Table S2). A similar pattern occurs in two additional regions of interest within the same feather (figure S5). Although the smaller proximal hooks in these sections are less distinct, this may reflect developmental variation during melanosome formation along the feather barb. Significantly smaller hooks were also observed in cross-sections of *in-situ*feather barbs from the anterior side of the feather crest (figure S6). Based on these observations, we propose that the hooked structures—particularly those on the dorsal, ventral, proximal, and distal sides of the melanosomes—enhance the structural integrity of the barb (figure S7). However, these features may be teratological and unique to this individual, as no similar structures have been reported in other species. These hooks may stabilize the stacked melanosome rods and contribute to increased barb dimensions, such as diameter and length. The sections exhibit modified (or asymmetric) hexagonally packed melanosomes with presence of extra hooked linkages (Figure 3c-d and e-f). The long rod-like melanosomes are different from all other known feather melanosomes from both extant and extinct taxa in having some extra hooks and an oblique ellipse shape in cross and longitudinal sections of individual melanosomes (Durrer 1986, Zhang, Kearns et al. 2010). The asymmetric packing of the melanosomes (the major axis leans leftward in cross section) played a major role in the reduction of fossilized keratinous matrix within the barbs, which may correspond to a novel structural coloration in this extinct bird. The close packed hexagonal melanosome pattern found in extant avian feathers yield rounded melanosome outlines in contrast to the oval-shaped melanosomes (see figure S8, x<y) in the perpendicular section here. The asymmetric compact hexagonal packing (ACHP) of the melanosomes is different from the known pattern of melanosomes formed in the structure of barbules among extant birds (Eliason and Shawkey 2012), which has been seen as a regular hexagonal organization. The packing of the melanosomes in an asymmetric pattern, on the microscopic level, might be related to the asymmetrical path of the barb extension direction observed at the macroscopic level (figure S5).

In the computational (FDTD) simulation, when light is incident at a small angle (<30°, both s- and p-polarized) in the X-Y plane, no evident reflection peak was found in the visible spectrum, as shown in Figure 4c (e.g., when a viewer observes at this angle, the light is almost completely absorbed, and the crest feather would have a darkened appearance). As the incident angle varies from 40° to 70°, the wavelength corresponding to the reflection peak shifts from 687 nm to 475 nm, and the color of the reflection peak gradually changes from red to deep blue (inset in Figure 4c clearly shows how the resulting color varies with the angle of incidence). From the perspective of optical modeling, the generation of iridescent color (reflection peaks in the visible spectrum) is caused by the constructive interference between scattered light produced by these uniquely shaped melanosomes that are compactly arranged in the barbs (Vinther, Briggs et al. 2010). Between blue and red, reflection peaks of green and yellow also are present, corresponding to the wavelengths of 512 nm and 578 nm respectively. It is worth noting that in the case of both p- and s-polarized incident light, the positions of the reflection peaks are nearly identical, with only a slight difference in peak reflectance. Therefore, the three-dimensional multilayer nanostructure model is somehow equivalent to one-dimensional multilayer biphotonic crystals (Saranathan, Forster et al. 2012). In summary, the crest feather of the Cretaceous enantiornithine shows a strikingly angle-dependent and iridescent coloration pattern. When we consider the shrinkage effect on melanosomes demonstrated experimentally (McNamara, Briggs et al. 2013) and various distances between melanosomes resulting from compression effects (Pan et al. 2019; Zhao et al. 2020), our additional FDTD modeling shows a similar reflection spectrum with a narrower bandwidth of the color peaks (Supplemental Materials).

**Figure 4.**
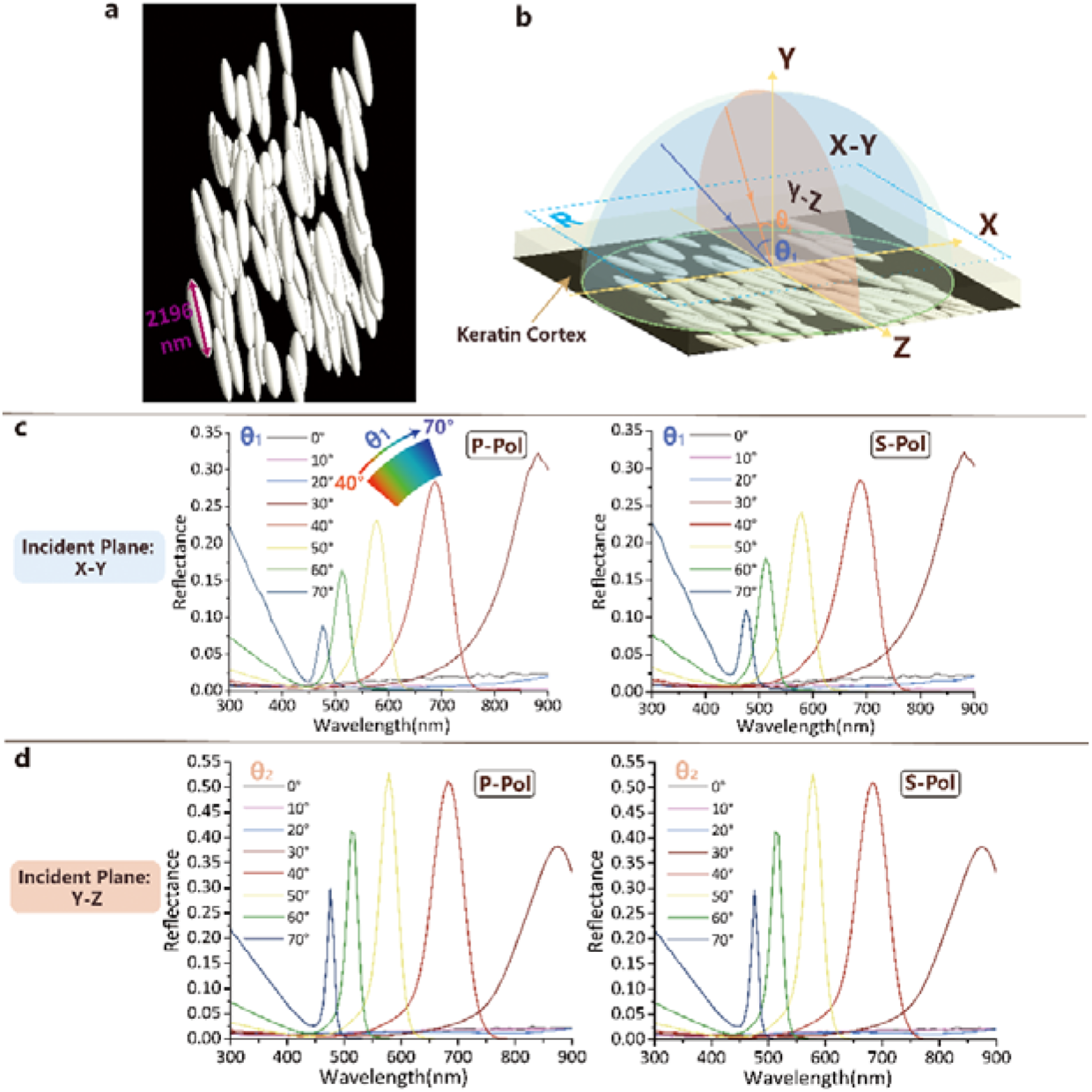
FDTD modeling results. (**a**) Representative multilayer melanosome model; (**b**) Schematic drawing of the reflectance calculation setup of the feather barb. X-Y incident plane (blue) is perpendicular, and the Y-Z incident plane (orange) is parallel to the special melanosome longitudinal axis. *θ*_1_ and *θ*_2_ are incident angles in X-Y and Y-Z incident planes, respectively. The blue dashed box (R) indicates the area where the reflection spectra are simulated and calculated. (**c**) Angle-dependent reflectance spectra calculated for p- and s-polarized light by FDTD modeling with X-Y incident plane. (**d**) Angle-dependent reflectance spectra calculated for p- and s-polarized light by FDTD modeling with Y-Z incident plane. Keratin cortex layer was considered in the modeling.

## Discussion

This study of a fossilized feather with a combination of microscopic imaging, ultrathin sectioning, and three-dimensional modeling has provided a more accurate examination of ancient structural coloration and uncovered unusual melanosome structures. Using previous conventional approaches alone, we would have classified this feather crest simply as black or close to iridescent based on its melanosome aspect ratio (Nordén, Eliason et al. 2021). The recognition of overall iridescence links to a commonly found extant bird color range only known previously from the platelet-shaped dinosaurian melanosomes (Hu, Clarke et al. 2018) and a few fossil feathers (Pan, Li et al. 2022) (Nordén, Faber et al. 2019, Nordén, Eliason et al. 2021). The independent evolution of overall iridescence, similar to that found in living birds, was reconstructed here based on a novel array of melanosomes in a feather barb in an extinct non-crown group bird (Nordén, Faber et al. 2019, Yoshioka and Akiyama 2021). The occurrence of iridescent structural coloration in an enlarged head crest on this ancient bird is similar to structures present across many living birds that employ iridescence for intimidation, signaling, and particularly for mate attraction or intraspecific territorial defense (Terrill and Shultz 2023). Given that this individual bird likely was a male based on the occurrence of paired rachis dominant tail feathers (figure S1), the vibrant coloration on a prominent feathered crest suggests that sexual selection may have long played an important role in avialan (feather) macroevolution (Foth 2020, Wang, O’Connor et al. 2021). The vibrant color and its presence on an enlarged feather crest likely are interlinked, and probably indicate an evolutionary and selective relationship among color, feather position, and feather morphology/size.

Our discovery of new melanosome shapes and networks suggests that birds evolved previously unrecognized patterns of color production outside of the limits present among crown group birds. Additionally, these pigments and their dense organization may have made the foundation of the feather darker and allowed for a more intense or vibrant iridescent coloration, a potential relationship that is understudied among living birds. While the interrelationship between melanin content in feathers and increases in structural strength have long been known (Burtt 1981, Burtt Jr and Ichida 2004), perhaps the increased density of melanosomes in this fossil feather barbs added even more structural strength to the feathered crest of this bird in compromise for its lack of a dominant rachis. Additionally, the interlocking hooks among the melanosomes might have aided individual feather barb in resisting shear forces. The modified closely packed melanosome arrays reconstructed here also are potentially advantageous in increasing barb structural integrity and stability in the long barb ramus, and likely affected the visualized spectrum during the display of iridescent coloration. We cannot completely rule out the increased melanosome density in the fossil feather is the result of taphonomy. Size reduction in melanosomes was found previously in a simulation experiment (McNamara, Briggs et al. 2013), but such a reduction is very unlikely to result in the highly organized and consistent melanosome packing present here (McNamara, Briggs et al. 2013, Zhao, Hu et al. 2020). To test the potential impact of melanosome shrinkage and potential taphonomic changes to their spacing, we conducted an additional analysis (See Supplementary Materials), and the results yield similar iridescent spectral peaks as the original setting.

Since the iridescent coloration of living birds is produced by the accumulation of melanosomes mostly positioned in barbules, the identified fossil barbs filled with melanosomes represents a unique phenotype related to an early feather evolutionary and developmental stage. Given the lack of barbules in most early primitive feather forms (or filamentary structures) in non-avialan dinosaurs and pterosaurs, the positioning of melanosomes in barbs might be a plesiomorphic feature for ornithodirans broadly (Xu 2019, Cincotta, Nicolaï et al. 2022). With different destinations of melanosome transport during feather formation, the pattern recovered here suggests fundamental differences in feather pigmentation among Cretaceous stem taxa as compared to crown birds. The melanosome-filled barbs evolved earlier in feather evolution before the later appearance of innovations such as the presence of a stiff rachis and colorful barbules observed across living feathers (Yu, Wu et al. 2002).

A focus on the two-dimensional examination of fossil feathers has uncovered a large number of black or dark melanosomes across many early birds and feathered theropods (Li, Clarke et al. 2014, Hu, Clarke et al. 2018). However, our three-dimensional study of a fossilized Jehol feather specimen demonstrates that ultra-histological detail and spatial relationships can be preserved at the microscopic level. Importantly, our discovery suggests that reexamination of previously published ‘black’ pigmented fossil feathers might uncover a wider occurrence of ‘surprising’ iridescence and structural coloration among these early birds and non-avialan taxa. Such discoveries will aid in documenting both independent and convergent evolutionary pathways in the history of birds. This research further demonstrates that Jehol fossil feather tissue can be treated as (microscopically) three dimensional specimens allowing for the collection of valuable comparative data and application of comparative analytical techniques that facilitate a better understanding of early feather evolution and function.

Here, we recovered iridescent coloration in fossil bird feathers that was produced by three-dimensional melanosome packing patterns. This discovery dramatically expands the range of fossil bird coloration space, and this diversity in feather coloration in an early bird fits well with its forested environment, as proposed by the coeval fossil plant community (Wu, Li et al. 2023). Tropical forest environments are accompanied by colorful flowing plants and green leaves, and the structural color of this feathered crest is advantageous for bird signaling as well. Forested environments (flowers, leaves, light, and shadow) can be advantageous for birds and their signaling as more colorful feathers are found today in tropical regions as demonstrated among extant vibrant coloration of passeriform birds in particular (Cooney, He et al. 2022).

## Materials and Methods

### Fossil material

The referred specimen of the enantiornithine bird *Shangyang* sp. (IVPP V 26899) was recovered near La-Ma-Dong Village in Jian-Chang County, within the Jiufotang Formation (dated ∼120 Ma). It can be referred to *Shangyang* by the presence of derived features in the premaxillae, tibiotarsus, and sternum (Figure 1 and Supplementary Materials).

The feather fragment (labeled sample Figure 1) extracted from the specimen were mainly analyzed using SEM (Scanning Electron Microscopy) and S/TEM (Scanning/Transmission Electron Microscopy). The extracted feather sample is one of the longest crest feathers, was displaced far from the caudal position of the skull (sample in Figure 1), and belongs to a fully mature asymmetric barb dominant feather. In addition to the singular feather, three fragments of other feathers were extracted from the *in-situ* position of the head crest and sampled to validate the revealed structure (Supplementary Materials, figure S6). A small fragment of the right femur was extracted to produce a ground-section to evaluate osteo-histological features in relation to its ontogenetic stage (figures S1 and S2).

### Scanning electron microscopy (SEM)

Feather samples coated with gold were examined with a FEI ESEM Quanta 450, which is operated at the Tectonic Laboratory in the China Geological Survey in Beijing, China. The SEM was operated with about 10 mm working distance of the sample under a high voltage of 20 KV. Not only the surface of feather sample was examined, but the raw material of the cross-section exposed in the ultrathin sliced sections also were imaged to characterize an overall distribution and thickness of soft tissue in comparison with the adjacent sediment or matrix (Figure 2 and figures S3). Additionally, a FIB-SEM (Heliox 5CX Due Beam system) also was further applied to validate the melanosome packing in an extracted volume, and the imaged block is about 11 x 13 x 9µm (figure S7).

### Scanning/Transmission Electron Microscopy

The crest feather fragments were embedded in epoxy resin using the SPI-PON 812 Embedding Kit (MNA, EPOK, DDSA, DMP-30)(Luft 1961) at the Key Laboratory of Vertebrate Evolution and Human Origins of the Chinese Academy of Sciences in Beijing in 2022. After a step-wised polymerization at 37°C for 12 hours, 45°C for 12 hours, and 60°C for 48 hours, the epoxy-embedded block was prepared and cut using an ultrathin slicing machine (Lecia EM UC7), equipped with a diamond knife (ultra 45°, 3 mm) for further TEM observations. The ultrathin slices were placed on copper grids with carbon film (200 meshes in ∼3 mm diameter copper gird). The thickness of a prepared ultrathin slice is approximately ∼70 nm, and the slices were consecutively placed on ∼3-5 copper grids. The removed thickness of the sample for making those ∼3-5 copper grids for one run of sectioning (serial numbered grids were made) took about ∼20 µm of the prepared pyramidal-cubic shaped block (Figures 3 and 5). The direction of slicing was either perpendicular to the long axis of the barbs, or nearly parallel with the long axis of the barb, for the two spots selected for cutting.

S/TEM imaging was conducted on a Thermo Scientific (FEI) Talos F200X G2 TEM at the IVPP, Beijing during 2021-2022. The Talos F200X was operated at 200 KV with an analytical scanning transmission electron microscope mode (S/TEM). TEM slices were made from the isolated feather adjacent to the head and rostral portion of the skull. The two specific micron-sized spots of the feather were targeted for examination of their microscopic features by ultrathin slicing and examination as shown in Figure 3. For the long contour feather, one spot close to the middle of the feather was made as a cross section of the barb, and for the other spot, the slice was made close to the left side of the fragment with a longitudinal slice (Figure 3). The strategic consideration of the two directions (and spots) of cutting was to examine the melanosomes and other features in different views in order to visualize these different features and obtain three-dimensional data (see Supplementary Materials).

### X-ray computed tomography

The fossil slab was scanned using a GE |phenix| micro-scanner with a resolution of 24.3 µm at Key Laboratory of Vertebrate Evolution and Human Origins, IVPP, Chinese Academy of Sciences, Beijing. For the skeletal anatomy of the bird, µ-CT scans and digital rendering were applied to aid in visualizing the skeletal features of the new specimen (Figure 1). The digital rendering of the specimen was completed using Avizo 9.0. (Figure 1). Measurements were made using digital calipers, as well as validated in the CT data digitally (Table S1). To achieve optimized image resolution, the field of view was adjusted, resulting in part of the feet being excluded from the CT scan.

### Technical protocol for ultrathin slicing

Prior to the work presented here, we obtained numerous practice sections from a few samples of fossil feathers. In our prior samples, the fragmentary feather tissues were used to improve the technician’s capability and precision in sectioning exercises by comparison with the desired direction and sectioned direction for given a fossil. There are several key steps elaborated below for the successful repetition of practices of sectioning in the correct alignment. First, we examined the fossil feathers with light microscopy and SEM prior to the embedding process to make sure the barbs were aligned correctly in the epoxy resin. Meanwhile, the small part of the extracted feather sample was aligned, using the microscope (Lecia UC 7), into a proper position during the emending and stage trimming process. The attention to feather direction was performed to assure that the barb major extension direction was aligned with the long axis of the bullet mold (embedding capsule). We also checked the position of the feather fragment in the SPI-Pon 812 during embedding and polymerization. The sliced section should be precise enough to acquire the desired images and information after planning and optimization of each step described above, and before the sectioning. Of course, the ultrathin slicing practices can improve the technician’s skill and increase cutting precision which require a gradual learning process.

### FDTD modeling of the feather’s reflectance

To explore the coloration of the three-dimensionally packed melanosomes, Lumerical finite-difference-time-domain (FDTD) software (Ansys Canada Ltd.) (Inan and Marshall 2011, Sullivan 2013) was used to simulate the visualization of light propagation on the crest feather with regular arrangements of melanosomes (Figure 1). The FDTD method allows explicit simulation of light interaction with particular-shaped melanosomes by numerically solving Maxwell’s equation in the time domain. The construction of a three-dimensional nanostructure model is based on our characterized data (Figure 4 and Table S2), and detailed modeling steps are included in Supplementary Materials. The dispersion of the complex refractive index that was applied in our simulation is described and shown in figure S9. In order to mimic real visual conditions, several variables were considered during the simulation, including angle of light incidence in two orthogonal incident planes (X-Y and Y-Z plane) (Figure 4b) and polarization state (s- and p-polarized). Usually, p-polarized light is understood to have an electric field direction parallel to the plane of incidence, and s-polarized light has the electric field oriented perpendicular to the incident plane. Reflectance spectra are presented in Figure 4c-d.

### Caveats about potential taphonomic melanosome changes

The simulated colors here are robust even when taking potential taphonomic changes into consideration. The chance that the highly organized and anatomically oriented nanoscale pattern formed here is the result of geological compression or pressure during fossilization and diagenesis is quite low, and such structures are absent or unseen among known taphonomic artifacts (McNamara, Briggs et al. 2013, Zhao, Hu et al. 2020). At present, we cannot exclude the possibility that sheer stress could result in the observed asymmetric packing of melanosomes. However, that pattern has not been reported in any known experiment or geological process of fossilization including those related to Jehol bird or non-avialan dinosaur fossils. Therefore, we suggest the pattern we obtained here represents original biological structures. Given experimental observations of melanosome shrinkage at high pressures and temperatures (∼10% shrinkage) (McNamara, Briggs et al. 2013, Zhao, Hu et al. 2020), we also completed additional modeling with the fossil melanosomes enlarged (110%, see Table S3) along with the fossilized keratinous cortex and matrix. This additional modeling resulted in all of the spectral peaks present in the original modeling being recovered. Therefore, our results are robust even when considering known potential taphonomic artifacts, and both approaches produce similar results with the red-to-blue iridescent coloration having been present in the crown feather (figure S10).

## Acknowledgments

We thank: Yang JM, Ma JB, Wen XL, and Li DS for technical assistance in sampling and ultrathin sectioning and STEM imaging; Hou YM and Yin PF for assistance with micro-CT scanning of the specimen; Gao W for photography; and Xu Y for the artistic reconstruction. Two reviewers are grateful in providing valuable suggestions for improving earlier version of the manuscript.

## Author contributions

Conceptualization: ZL and ZZ

Methodology: JH, MY and ZL

Investigation: ZL, TAS, JH, YM, and JAC

Visualization: JH and ZL

Supervision: ZL and ZZ

Writing—original draft: TAS, ZL, YP, MW, JL and TZ

Writing—review & editing: ZL, JH, TAS, MY, MW, YP, TZ, JL, ZZ, and JAC

## Data and materials availability

All data are available in the main text or the supplementary materials

## Supplementary Materials

