## Supplemental file for "Iridescent structural coloration in a crested Cretaceous enantiornithine bird from Jehol Biota"

Li Zhiheng *et al.*

**This PDF file includes:**

Supplementary Text

Figs. S1 to S10

Tables S1 to S4

Supplemental References

Supplementary Text

1. Systematic Paleontology

Aves Linnaeus, 1758

Pygostylia Chiappe, 2022

Ornithothoraces Chiappe, 1995

Enantiornithes Walker, 1981

Genus *Shangyang* Wang et al. 2019

1.1. Brief description and diagnosis of *Shangyang* specimen

The small enantiornithine fossil specimen (IVPP V26899) described here can be referred to *Shangyang* sp. based on the derived skeletal anatomy shared with the holotype (IVPP V25033). For example, the presence of fused premaxillae along their entire length, a nearly square sternal outline, and the elongate fibula help to diagnose the fossil (Wang and Zhou 2019).

The intermetacarpal space between the major and minor metacarpals is quite narrow (Figure 1 and figure S1). The minor metacarpal is slightly curved and extends distal to the major metacarpal. The main part of the sternum has a square outline, and the lateral trabecula extends caudally, expanding into a large triangular plate (Figure 1). The midline projection of the sternum is stout and extends about the same length as the lateral trabecular process (Figure 1). The fibula is long, being over two thirds of the length of the tibiotarsus. The distal pubis is expanded into a boot shape at its distal end (figure S1).

Differing slightly from the holotype, the furcula of IVPP V26899 has a longer hypocleidium. The craniolateral process of its sternum also protrudes less than the holotype, and that might relate to its sub-adult ontogenetic stage based on histological examination of its leg bones. In addition, the size of the newly referred specimen is about 12 cm measured from head to tail, which is close to the size of the holotype (Figure 1).

1.2. Locality and Horizon

The new fossil specimen was recovered from La-Ma-Dong Village in Jian-Chang County in western Liaoning Province, northeastern China. Based on previous geological research, the fossil bird derives from the Lower Cretaceous Jiufotang Formation with the fossil-bearing strata dated as approximately 120 Ma (Yu, Wang et al. 2021).

1.3. Summary of enantiornithine feathers including the new specimen

Enantiornithes are the most diverse avialan clade among Mesozoic birds (Chiappe and Walker 2002) with particularly abundant and spectacularly preserved fossil specimens from the Early Cretaceous Jehol lagerstätte in northeastern China (Zhou, Barrett et al. 2003, Zhou and Zhang 2007). Those enantiornithine fossils from the Jehol Group preserve a surprising variety of morphological disparity and diversity not only in their skeletons, but also in their striking plumages across their limbs, heads, and tails (Wang and Zhou 2017). Enantiornithine feather assemblages are represented not only by the densely distributed contour and down feathers that covered their whole bodies that functioned largely in insulation, but also by atypical feather morphologies on their heads and other modifications like elongated rectrices (Foth and Rauhut 2020). This morphological diversity across enantiornithine plumages suggests that they played a major role in display, signaling, and sexual selection (Wang, O’Connor et al. 2021). Those hypothesized functions are reinforced by the preservation of ancient melanosome pigments in the microstructure of the fossil feathers that would have resulted in a variety of colors, demonstrating the long evolutionary history of feather coloration in the clade.

The well-developed preserved integumental structures of IVPP V26899 are mainly pigmented in a currently black-brown color within the white-grey matrix, distributed surrounding the whole body (Figure 1). This plumage not only includes elongate pennaceous flight feathers associated with its forearm, but also a dense covering of contour feathers (including down feathers) distributed along the forehead, cranium, neck, and caudal vertebral region (Figure 2). The feathered crest attached to the forehead and cranium of the new specimen is quite darkly pigmented (Figure 1). The erect feather bundles show a striking pattern with gradually increasing shaft (or barb) length towards the caudal end of the skull. The highly projected contour feathers approach the occipital region of the skull, forming a triangular crest (Figure 1). In addition, the shallow impressions of the paired rectrices are ribbon-like, and associated with the pygostyle (figure S1). The long pennaceous flight feathers (or remiges) are well developed and articulated with the forelimbs, including long primaries and secondaries (Figure 1).

The long and well-developed feather crest covers most of the dorsal surface of the skull (Figure 1). The crown (or crest) feather barbs (or shaft) gradually increase in length and reach their maximum lengths near the caudal part of skull. The shaft of the longest crest feather is about 2 cm, even longer than the dorsoventral height of the skull (Figure 1). The overall shape of the crown feathers forms a triangular crest that is projected dorsally and caudally. The rostral end of the feathered tract extends rostrally down the forehead along the premaxillae (figure S1). The crest feathers on the caudal side are longer than the rostral ones. They are highly projected on the parietal region and a silt-like fissure appears to be present between the feathered bundles attached to the frontal and the parietal (figure S1).

The long contour feathers extending from the caudal part of the head are composed of multiple slightly curved barbs, shortly branching off from the calamus at its base (figure S1). Densely packed clusters that comprised of elongate oval-shaped melanosomes are visible in the surface and the crack region of the feather barbs when examined with the SEM (Figure 2).

1.4. Description of Skeletal anatomy

The entire specimen (IVPP V26899) was compressed largely dorsoventrally, except for the laterally preserved skull (Figure S1). The skull is proportionally long, exceeding the length of the humerus. There are three or four conical-shaped teeth present on the premaxillae, and six teeth on the maxillae. At least eight alveolar pits are recognized on the dentary bone, indicating missing teeth. The dorsal surfaces of premaxillae are straight, and a T-shaped lacrimal bone is present. The external narial opening is silt-like, and the antorbital fenestra has a triangular outline. The frontal bones are crushed laterally, with the left and right sides appearing to be fused (figure S1).

There are at least eight or nine cervical vertebrae, and they are all exposed in ventral view (figure S1). The lateral edge of the coracoid is expanded slightly. The length of the hypocleidium is about the same length as that of the furcular rami. The lateral trabecula of the sternum is caudally expanded into a triangular process. The humerus has a dorsal supracondylar process. The alular metacarpal (cmI) is fused with the major and minor metacarpals proximally, but distally the two longer metacarpals are separate. The proximal part of the left scapula appears to be broken and incompletely fused during healing from a wound (figure S1). The length of the first digit is about the half of the length of the carpometacarpus. The total length of the forelimb (humerus + carpometacarpus + manual digit) is shorter than that of the hindlimb (Table S1).

The distal end of the fused pubis bares a booted process. The pygostyle is distally tapered, forming a spear shape (figure S1). The ilium has a dorsal bulge near its middle portion. The fibula is comparatively longer than that of other enantiornithines, reaching over two thirds of the tibiotarsus length. Proximally, metatarsals II-IV are fused as in other enantiornithines. Metatarsal III is the longest among metatarsals II-IV, with metatarsal IV slightly shorter than metatarsal II. The ungual (claw) size is large in all four pedal digits (figure S1), and the ungual lengths in digits I and II are even longer than those of the respective penultimate phalanges.

Ground sections were acquired from the right side of the femur to assess the osteo-histological features of the bone and its ontogenetic stage. As shown in figure S2, long, flat-shaped lacunae are present widely and densely packed throughout the major part of the bone section. Very few secondary osteocytes are present, and parallel-fibered bone tissue is underdeveloped. The flattened osteocyte lacunae dominate the cellular shape, with observable vascular canals connecting different lacunae. Overall, the osteo-histology indicates that the bird was still in an active growth stage at the time of death, suggesting it was in its sub-adult growth phase.

2. Microscopic methods and ultrastructural morphology of the feather tissue

2.1. Ultrathin Sections: Preparation and Results

We examined two different micron-sized spots for the feather fragment extracted from the new specimen IVPP V26899 (Figures 1-3). The surface size of the prepared cubic block in top view for cross section slicing is about 170 $\times$120 µm (figure S3).

The advantages of our multiple selections for targeted sampling adopted here are demonstrated by uncovering different aspects of feather and melanosomes ultrastructure. In addition, our consecutive ultrathin sectioning of one spot removed a tissue thickness of approximately 20 μm, which was measured from the lower to upper bound of those slices in a given single position. All of those slices were picked up consecutively with 3-5 serially numbered copper grids with carbon film.

High-resolution TEM/STEM (Transmission Electron Microscope/Scanning Transmission Electron Microscopy) imaging of the well-selected spots of sampled feathers were used to investigate the ultrastructure regarding melanosome geometry and keratinous tissues in the thin sections in different views. Our new microscopic data strongly suggests that soft tissue can be preserved well, and it was revealed further in TEM examination. The detailed features revealed here indicate an exceptional preservation of three-dimensional soft tissues in the slab fossil. Our measurements of thickness of the feather tissue measured from the top view of the section block are about 10-30 µm, figure S4 (b-d).

2.2. Crest feather samples

The long crest feather was sectioned in two directions with one series made on the side of a barb longitudinally and another series made across the cross section of the central barb (Figure 3). The three-dimensional (3-D) packing of the melanosomes was discerned in both the cross- and longitudinal- views respectively, with full details (Figure 3). The unique asymmetric-hexagonal-packing (AHP) discovered here defines a dense stack of melanosome assemblage array, with additional hooked linkages among the neighboring melanosomes (Figure 3). The novel shapes and packing of the melanosomes played a major role in the reduction of air or keratinous space within the barb, and this may correspond to the unique type of structural coloration modeled below. The tiny hooks identified here in some areas, as extra connections, linked the ‘head’ and ‘tail’ portions of the sausage-like melanosomes (Figure 3). They may have increased further the abrasion resistance of the barbs. The modified sausage-like melanosomes with novel packing have not been reported from any feathers of extant or extinct birds (Durrer 1986, Zhang, Kearns et al. 2010). Differing from the normal compact hexagonal packing (Durrer 1986), the arrangement of these melanosomes appears to be rotated toward the right-lateral side of the feather barb, major axis of the melanosomes lean left-ward (figure S1). This novel packing could be acquired in the enantiornithine feather through a modification of the simple rodlet-shaped melanosomes previously reported (Zhang, Kearns et al. 2010).

In addition to the middle portion, we sectioned two additional positions from more proximal sides of the feather barbs (figure S5) to validate the melanosome packing. As expected, very dense melanosomes were observed. The hooked structures are less clearly defined because of their smaller size or developmental immaturity. However, the asymmetric packing is similarly present, with the melanosome on the right side packed over the melanosome on the left (i.e., longer axis leans toward left in cross section view). As we argued in the main text, this structure in melanosome hooks may help increase the dimension of the barb size, allowing it to grow thicker and enhancing the structural integrity of the barb. The same pattern also is observed in the cranial barbs sampled from the anterior crest feather, which has a much shorter length at the front of the head (see figure S6).

2.3. FIB-SEM 3D Tomography

The milled block was extracted from the central portion of the long crest feather sample using a Helios 5 CX Dual-Beam system at the IVPP. Before extraction, the surface of the targeted area of fossil sample was deposited with carbon and tungsten (GIS port) and then milled using gallium ions. The nanomanipulator (Easylift) attached to the sampled block for lift-out and attached to a TEM copper grid for further imaging. The sample was milled 20 nm for each time along the ‘z’ axis and imaged serially (figure S7). Resulting serial images revealed intricate structures, such as linkages and hooks, within the stacks of melanosomes.

2.4. Identification of barbs

The overall morphology of the feather represents a common type of primitive feather with several barbs diverging just above the calamus. The major barbs in the middle appear to be slightly larger and thicker in diameter than the neighboring barbs. No evidence of a rachis or barbules was found in the cross section, meaning that the branched barbs are the major components of the feather tissue preserved. The largest diameter of a barb appears to be over 10 µm, exceeding the diameter of most barbules in living feathers (figure S3). Given the overall evidence of the morphology and microscopic data (figures S3 and S4), we interpret the feather as being an asymmetric downy barb-composed feather, without a stiff central shaft and distal branched barbules. It is composed only of several laterally diverged barbs along the same direction with a fusion near a short calamus at its base (figure S4).

3. FDTD modeling of the crown feather reflectance

Both cross- and longitudinal-section anatomy of the area of crown feather with particular-shaped melanosomes are shown in figure S8a-b. To simulate the spatial and spectral responses of the crown feather, the three-dimensional FDTD modeling is utilized based on the anatomy results aforementioned (figure S8a-b). Additionally, the numerical simulation results related to the angle-, polarization- and wavelength-dependent reflectance spectrum were obtained. In the simulation, novel hexagonal-packed particular-shaped melanosomes were modeled, and in particular, hexagonally arranged oval-shaped (ellipsoidal) microstructures were employed to simulate the derived arrangement of sausage-like melanosomes as shown in figure S8c. The crown feather model consists of multiple layers of particular-shaped melanosomes and keratin wrapped around them (10 layers, each layer contains 10 melanosomes) as shown in figure S8d. The lengths of semi-axes of the triaxial ellipsoid are assigned according to the statistics of the SEM images (Table S2). The period of hexagonal arrangement (e.g., twice of side length of the hexagon) is selected as 406 nm to achieve similar compactness of the model and SEM images (Table S2). Specifically, the ellipsoidal melanosomes are arranged in the simulation region so that some of the adjacent melanosomes were in contact with each other to simulate the hooked linkages as shown in Figure 3d. The incident light was set accordingly to the s- and p-polarized of two orthogonal incident planes (X-Y and Y-Z plane), respectively. It penetrates through all of the melanosomes, undergoes near-field interactions (such as scattering, interference, etc.), and the reflected light is collected by the reflectance monitor above the incident light plane.

Melanosome (or melanin) is a strongly absorbing pigment, and hence has a complex refractive index (
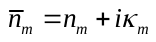
), in which the real component corresponds to the familiar refractive index of non-absorbing matter and the imaginary part accounts for light absorption (Wilts 2013). For all simulations, we used published values (Leertouwer, Wilts et al. 2011, Eliason and Shawkey 2012, Eliason and Shawkey 2012, Wilts 2013, Stavenga, Leertouwer et al. 2015, Jiang, Wang et al. 2018) for the refractive indices of keratin ($n_{k}$) and special melanosomes ($n_{m}$). The wavelength dependence of the real part and imaginary part of refractive indices are described by Cauchy equation and exponential function as follows:

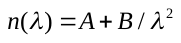
(1)

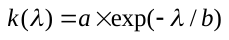
(2)

For concentrated melanin
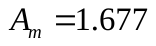
and
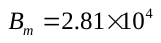
nm^2^, as for extinction coefficient of melanin,
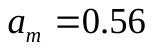
and
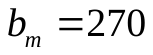
nm. In addition, for real part of the refractive index of keratin,
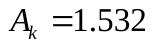
and
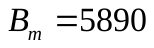
nm^2^, in all simulations, and the imaginary component of keratin is assumed to be negligible (Wilts, 2013). In summary, the dispersion of complex refractive index utilized in our simulation is described in figure S9.

Non-uniform mesh grids were utilized to obtain a trade-off between sufficient simulation accuracy and simulation time. In addition, we modeled only parts of the crown feather (one hundred novel-packed melanosomes with particular-shaped) to reduce computational resources. Stabilized and perfect matched layers (stabilized PMLs) are applied in all directions to avoid boundary reflection. The simulations were performed on a workstation. The processor of the workstation is two Intel Xeon Platinum 8280; one of each containing 28 cores and 56 threads, and each core has a base frequency of 2.70 GHz. For a single simulation running (e.g., computation for single wavelength, one polarization state (s- or p-polarized), and one incident angle in particular incident plane (X-Y or Y-Z) combination) takes the memory of approximately 130 GB, and the simulation time of about 12 hours.

4. Additional FDTD modeling

Taking into consideration the effect of melanosome shrinkage observed in feather melanosome taphonomic experiment under various temperatures and pressures, we performed additional FDTD simulations to account for the melanosome size change (enlarged melanosomes used as shown in Table S3). The simulated reflectance spectra and comparison with the original simulation are shown in figure S10 and Table S4. It should be noted that the wavelength of the reflection peak in the 10% increased melanosome size setting (compare the original measurements) while considering taphonomic change of the melanosomes is almost identical to the original simulation under different incident light. However, the bandwidth of the reflectance peak decreased dramatically (i.e., the reflectance peaks look sharper), which means the feather with enlarged melanosomes should look more strikingly colorful than the normal ones.

**Supplemental figures and tables**

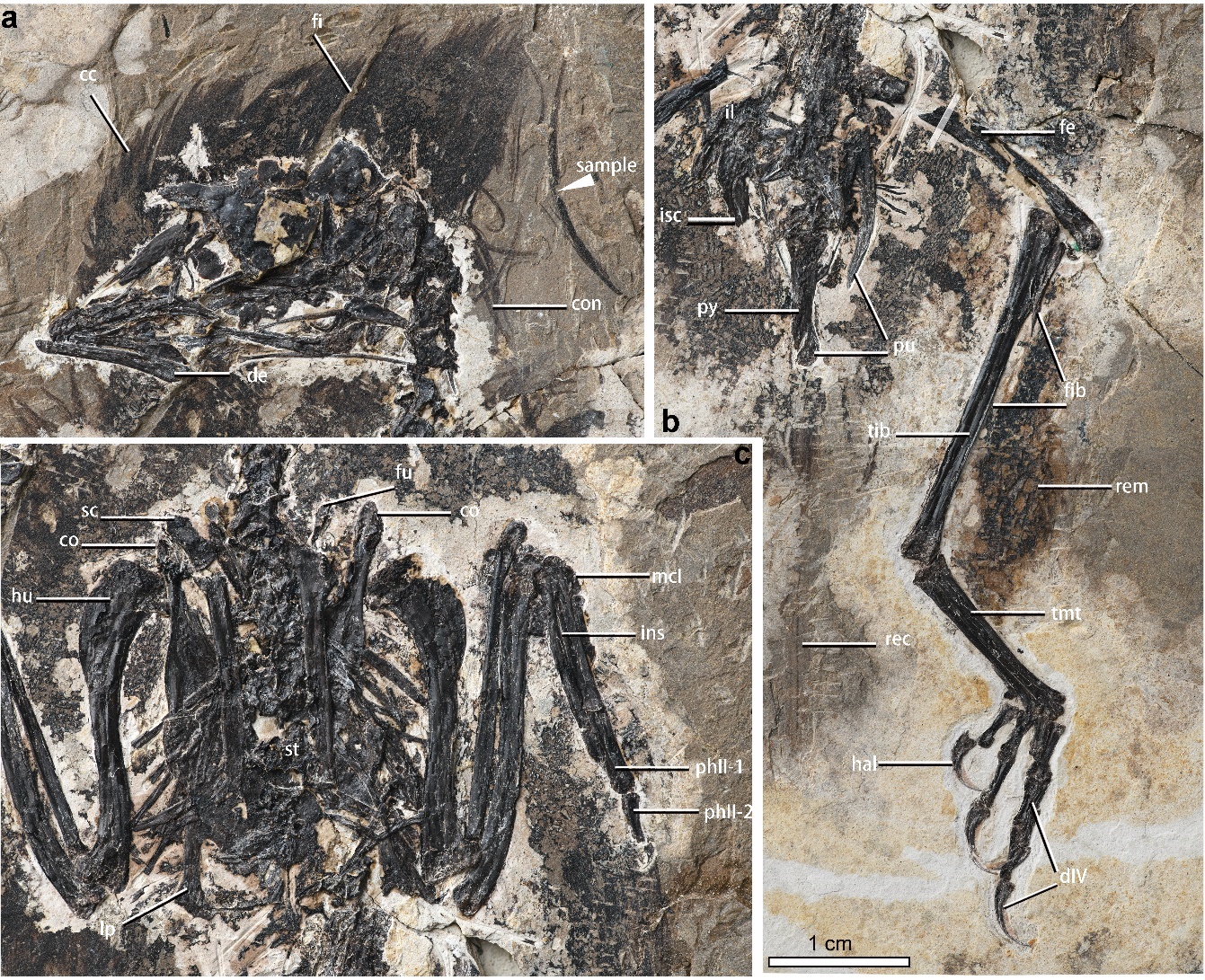

**figure S1**. Photograph showing details of skeletal and integumentary anatomy of the new enantiornithine specimen (IVPP V26899): (a)-skull, (b)-pelvis and hindlimb, and (c)-pectoral region. The sectioned feather is indicated as sample. Abbreviations: cc, cranial crest; co, coracoid; con, contour feather; de, dentary; dIV, pedal digit IV; fe, femur; fi, fissure; fib, fibula; fu, furcula; hal, hallucal ungual; hu, humerus; il, ilium; isc, ischium; lp, lateral process; mcI, metacarpal I; pu, pubis; py, pygostyle; rec, rectrices; sc, scapula; st, sternum; tib, tibiotarsus; and tmt, tarsometatarsus. White transparent bar in (b) indicates the region of the sampled femur fragment for ground sectioning.

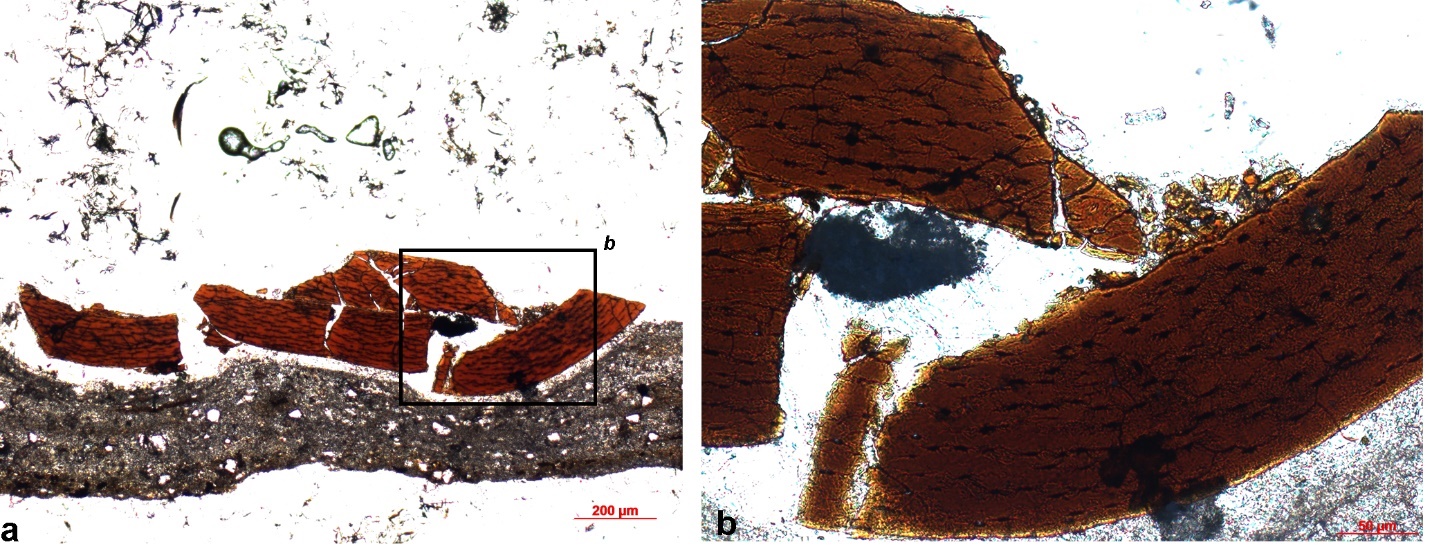

**figure S2. Microscopic ground section images to show the osteo-histological features of the femur sampled (a)-overall view; (b) enlarged view labeled in black box.**

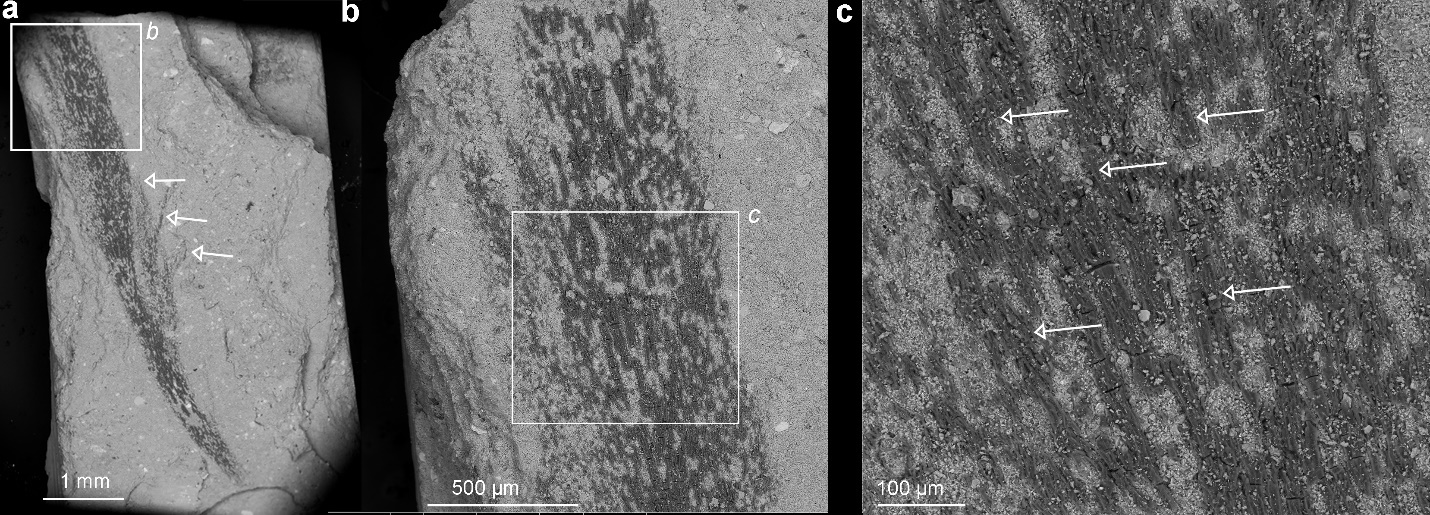

figure S3. SEM of the serial zoomed view (a to c) to show the lower portion of the sampled crest feather below the sectioned fragment. The individual barbs are roughly identified and labeled with white arrows; the feather tissue is preserved in stark contrast to the rock matrix under BSD (back scatting diffraction) model in the SEM imaging.

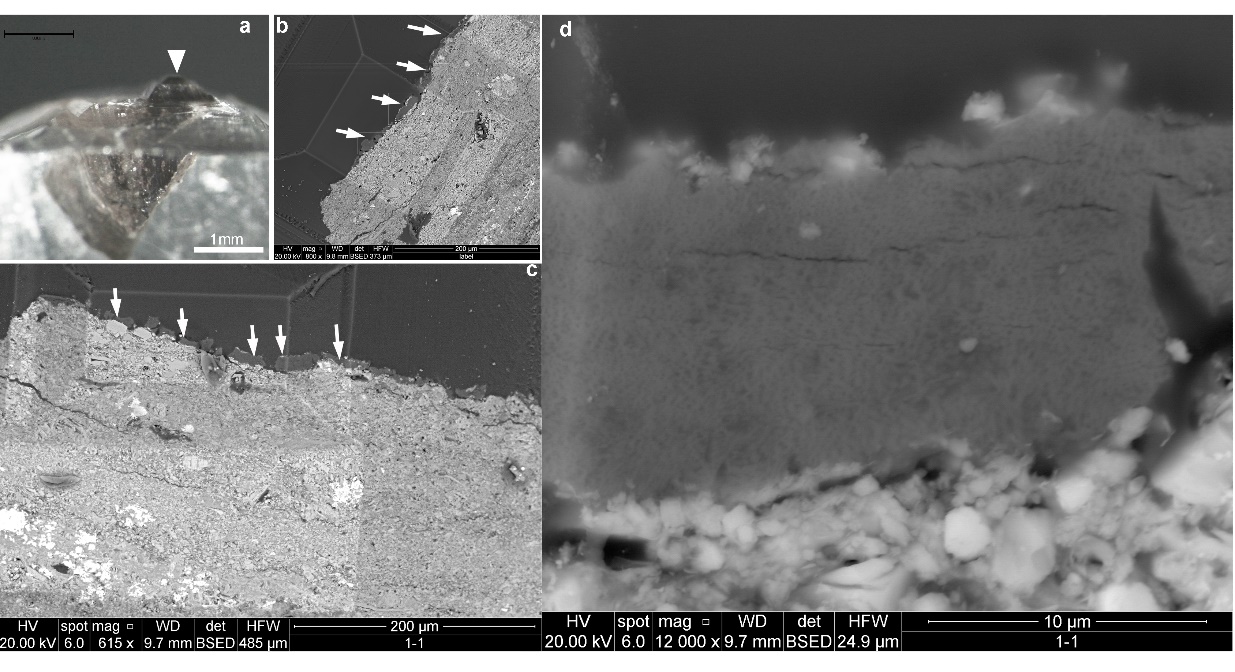

figure S4. Barb surface and cross section morphology of the sampled fragment of the crest feather. (a)-light microscopy; (b, c, d) - SEM images of the cross-section to show the melanosome-composed barbs in the sectioned region of the feather fragment. The white arrow in a indicates the trimmed stage for cross sectioning; the arrows in b and c indicate the individual barbs with space in-between; the zoomed image (d) shows the layer of melanosomes which entirely comprised the internal region of one barb.

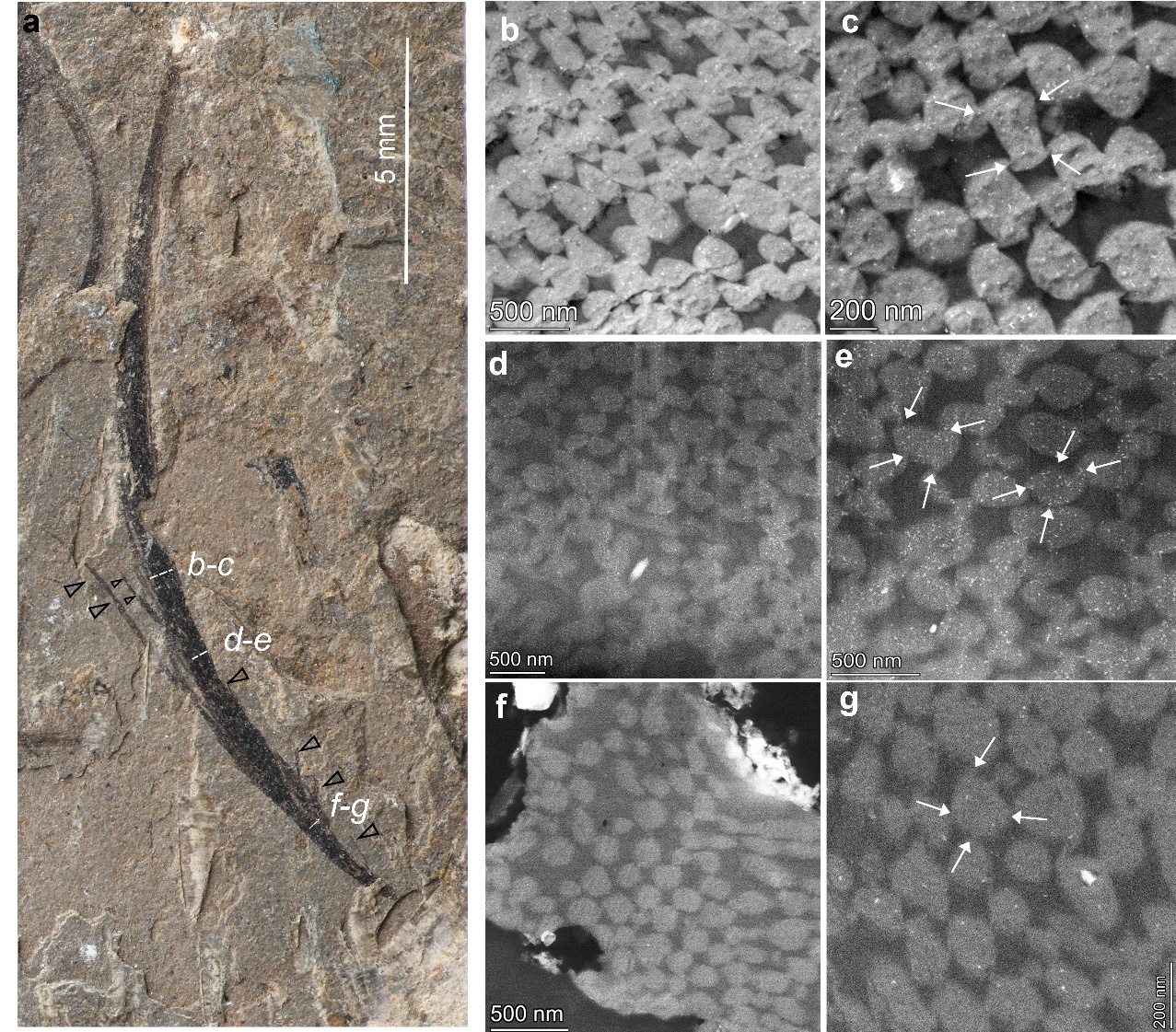

**figure S5. STEM images of cross-sections taken from three different positions (indicated by white dashed lines in a) demonstrate similar melanosome packing styles.** Dashed-lines labeled in (a) indicate where the corresponding position of these sections were taken, black arrows indicate the individual barbs that accumulated together in this long crest father. One distinct feature of these sections is the hooked-link structure that aligns the melanosomes into a modified hexagonal, packed arrangement. White arrows (in c, e, g) indicate the hooked structures observed in the selected melanosomes.

**figure S6. STEM images showing melanosome structure from three fragments of the feather crest (indicated by dashed lines and a white box in a) reveal the hooked linkages between melanosomes and their surrounding melanosomes structures in (b), (c) and (d).** Given the shorter length of these feather barbs, the hook structures are not as well-defined as those in the longer feather samples shown in the main text.

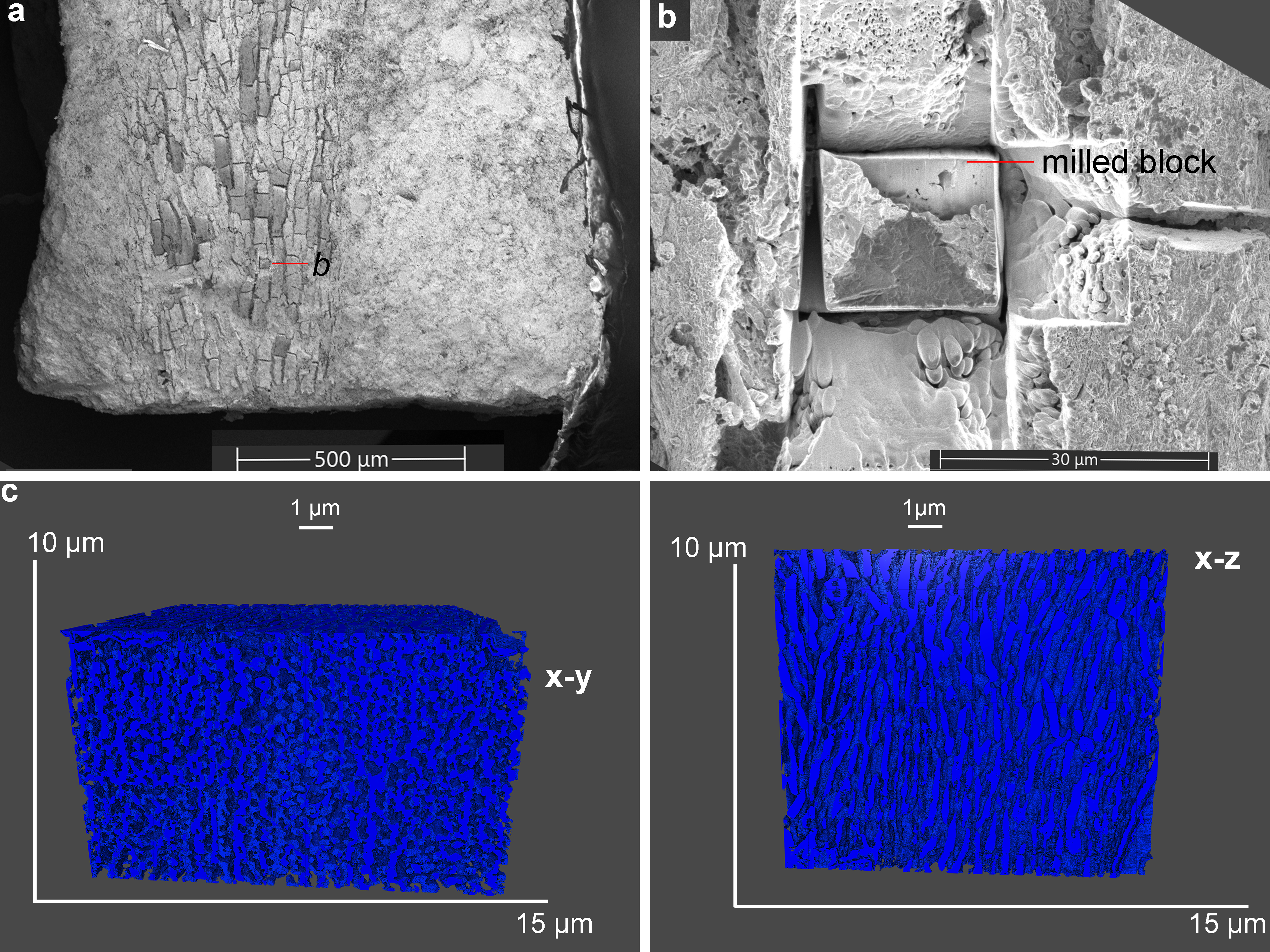

**figure S7. Targeted feather barb block prepared in FIB-SEM, with volume rendering reconstruction based on the acquired sequential cross-sectional images;** the volume reconstruction is visualized in the x-y plane (c-cross section view) and in x-z plane (d-sagittal section view).

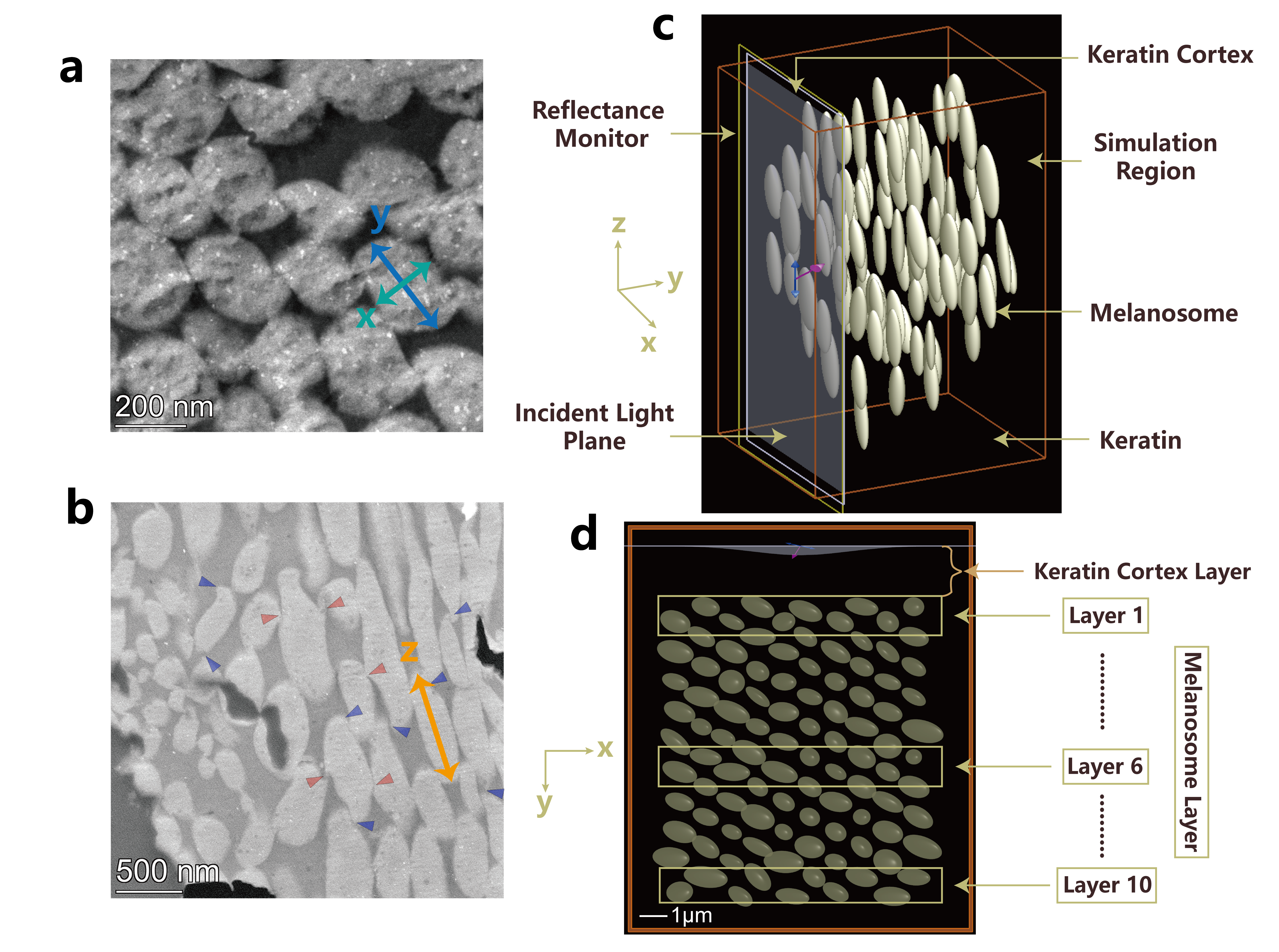

**figure S8. STEM images obtained from feather barb slices with the reconstructed 3D model melanosomes sets.** (a) Cross-sectional TEM image obtained from feather slice. x and y represent the length of minor and major principal axes of melanosome ellipsoid. (**b**) Longitudinal section TEM image. **z** represents the length of longitudinal axes of the melanosome ellipsoid. (**c**) Snapshots of the FDTD modeling for the multilayer crown feather. (**d**) Multilayer melanosome structures of the crown feather within the XY cross-section of the FDTD modeling. The thickness of cortex layer modeled is about 1.8 μm.

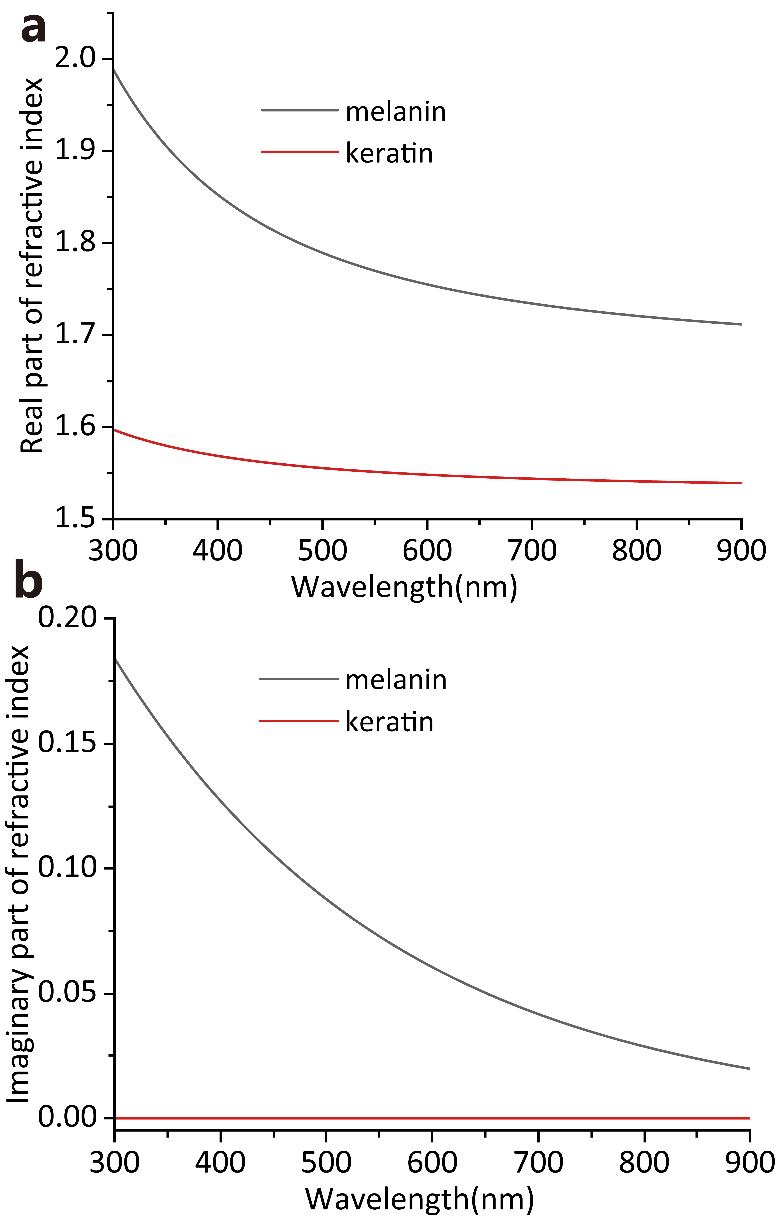

**figure S9.** **Refractive index dispersion.** The complex and wavelength-dependent refractive index ($\bar{n}=n+i\kappa$) for melanin and keratin utilized in our FDTD modeling of the barb reflectance spectra. The dispersion of the real part of refractive index **(a)** and imaginary part (or extinction coefficient) **(b)** are shown.

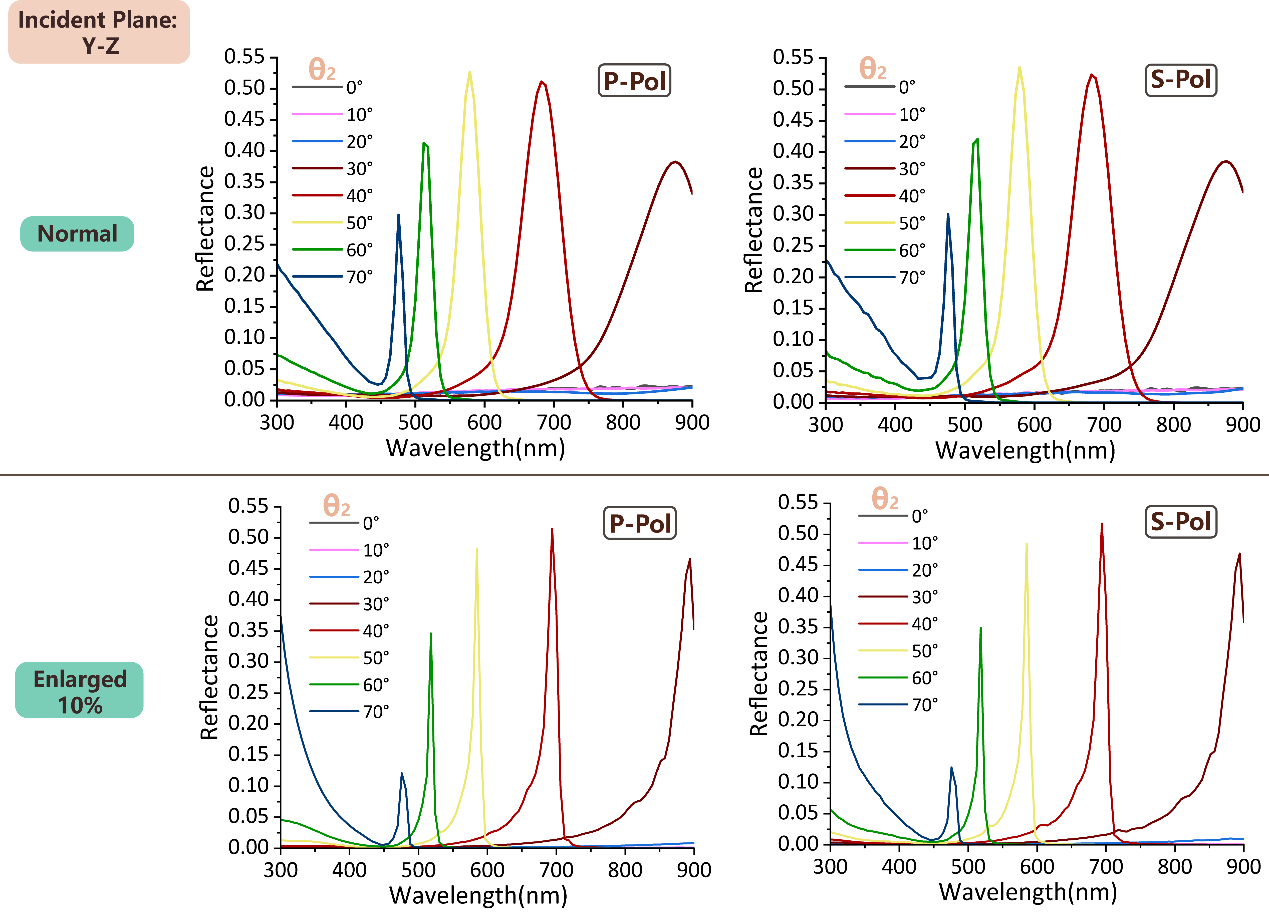

**figure S10. Comparison of angle-dependent reflectance spectra in the Y-Z incident plane for the normal and enlarged cases with consideration of melanosome shrinkage in taphonomy.**

Table S1. Measurements of major bone elements in IVPP V26899 (in mm).

| Skull length |  | 30.8 |  |  |
| --- | --- | --- | --- | --- |
| Humerus length | left | 22.6 | right | 22.2 |
| Ulna length | left | 23.6 | right | 24.4 |
| Scapula length | left | 19.2 |  |  |
| Radius | length | 23.6 | right | 23.3 |
| Sternum length | midline | 13.8 |  |  |
| Femur length | left | 21.2 |  |  |
| Pygostyle | length | 8.5 |  |  |
| Metacarpal II | length | 11.5 |  |  |
| Ilium | length | 13.1 |  |  |
| Pubis length | length | 17.2 |  |  |
| Ischium length | length | 6.5 |  |  |
| Metatarsal III | left | 15.0 | right | 15.5 |
| Tibiotarsus length | left | 27.0 | right | 27.5 |

Table S2. Measurements of the hexagonal-packed ellipsoidal melanosome modeling obtained from partial cross- and longitudinal STEM images (see figure S8a-b)

| Parameters | Mean | Standard deviation | Hexagonal period |
| --- | --- | --- | --- |
| Length (x) | 186.17 nm | 33.53 nm | 406 nm |
| Length (y) | 283.49 nm | 46.44 nm |  |
| Length (z) | 1774.03 nm | 269.21 nm |  |

**Table S3 Melanosome modeling employed in additional FDTD simulation (enlarged 10% compared to Table S2)**

| Parameters | Mean | Standard deviation | Hexagonal period |
| --- | --- | --- | --- |
| Length (x) | 206.65 nm | 37.19 nm | 446 nm |
| Length (y) | 314.67 nm | 51.55 nm |  |
| Length (z) | 1969.14 nm | 298.62 nm |  |

**Table S4 Reflectance peak wavelengths at different incident angles for the normal and enlarged cases**

| Incident angle (degree) | Reflectance peak wavelength  in original sets (nm) | Reflectance peak wavelength at 110% sized case (nm) |
| --- | --- | --- |
| 30 | 875.76 | 893.94 |
| 40 | 681.82 | 693.94 |
| 50 | 578.79 | 584.85 |
| 60 | 518.18 | 519.20 |
| 70 | 475.76 | 475.60 |
